# Reward prediction errors create event boundaries in memory

**DOI:** 10.1101/725440

**Authors:** Nina Rouhani, Kenneth A. Norman, Yael Niv, Aaron M. Bornstein

## Abstract

We remember when things change. Particularly salient are experiences where there is a change in rewards, eliciting reward prediction errors (RPEs). This feature of memory may be useful because it can help us find greater rewards and avoid lesser ones in the future. How do RPEs influence our memory of those experiences? One idea is that this signal directly enhances the encoding of memory. Another, not mutually exclusive, idea is that the RPE signals a deeper change in the environment, and leads to the mnemonic separation of subsequent experiences from what came before, thereby creating a new latent context and a more separate memory trace. We tested this in four experiments in which participants learned to predict rewards associated with a series of images within visually-distinct “rooms.” High magnitude RPEs indicated a change in the underlying distribution of rewards. To test whether these large RPEs created a new latent context, we first assessed recognition priming for sequential pairs that contained or did not contain a high-RPE event, as well as out-of-sequence pairs (Exp. 1: n=27 & Exp. 2: n=83). We found evidence of recognition priming for both sequential pair types, including the pair with the high-RPE event, indicating that the high-RPE event is bound to its predecessor in memory. Given that high-RPE events are themselves preferentially remembered (Rouhani et al, 2018), we next tested recognition priming for pairs that had one item in between them (i.e. the pairs were either across a high-RPE event or not), where none of the tested items were high-RPE items (Exp. 3: n=85). Here, sequential pairs across a high-RPE no longer showed recognition priming whereas pairs within the same latent reward state did, providing initial evidence for an RPE-modulated event boundary. We then investigated whether RPE event boundaries disrupt temporal memory of those events (Exp. 4). After reward learning, we asked participants to order and estimate the distance between two events that had either included a high-RPE event between them, or not. We found (n=49) and replicated (n=77) worse sequence memory for events across a high-RPE event. Altogether, these findings demonstrate that high-RPE events are both more strongly encoded and act as event boundaries that interrupt the sequential integration of events. We captured these effects in a variant of the Context Maintenance and Retrieval model (CMR; Polyn, Norman & Kahana, 2009), modified to incorporate RPEs into the encoding process.

## 1. Introduction

A single experience can change our expectations of future rewards. The ability to infer this change is critical to adaptive behavior, as it guides decisions to seek or avoid that experience in the future. For example, imagine you watch a new episode of what had long been your favorite television show, only to find that you strongly dislike it. Worse, this bad episode indicates a decrease in the show’s quality (e.g. brought on by a change in writers). In reinforcement learning, a surprising event (e.g., a dramatically substandard tv episode) generates a large reward prediction error (RPE), which quantifies the difference between expected and received reward. Recent work shows that larger positive or negative RPEs experienced during reward learning lead to improved memory for those surprising events (Rouhani, Norman, & Niv, 2018). However, the mechanism behind this enhanced memory is unclear. Is the episode where the quality of the show changed better remembered because it is more strongly stamped in memory? Or is it better remembered because it predicts a meaningful change in the state of the show, thereby separating the pleasant episodes that came before it from the unpleasant episodes that followed, creating separate clusters in memory? In other words, do high RPEs lead to better memory because they bind events more rigidly to the context in which the event occurred, or because they lead to the creation of a new context?

If high RPEs create a new latent state or context, then we predicted they would act as *event boundaries* in memory. In fact, prediction errors (outside of the reward domain) are thought to create event boundaries by segmenting the continuous stream of experience into separate memory traces (DuBrow, Rouhani, Niv, & Norman, 2017; Gershman, Radulescu, Norman, & Niv, 2014; Zacks, Speer, Swallow, Braver, & Reynolds, 2007). It is, however, unknown whether changes in the distribution of rewards, signaled by high RPEs, act as event boundaries in memory. Events boundaries structure the temporal organization of memories by interrupting the integration of events across them. This leads to worse memory for the order of events (“sequence memory”) and greater perceived distance for events across rather than within contexts (Dubrow & Davachi, 2013; Horner, Bisby, Wang, Bogus, & Burgess, 2016). This is further predicted by greater representational dissimilarity of those events in the hippocampus (DuBrow & Davachi, 2014; Ezzyat & Davachi, 2014). Interestingly, like high-RPE memories, memory for the event boundary itself is enhanced (Heusser, Ezzyat, Shiff, & Davachi, 2018; Swallow, Zacks, & Abrams, 2009). However, temporal memory for the events across the boundary is worse, suggesting a trade-off between memory for the boundary event and the mnemonic integration of events across the boundary (Heusser et al., 2018).

In four experiments, we investigated whether latent shifts in the reward distribution of a Pavlovian reinforcement task (which generate high RPEs) create such event boundaries in memory. In all experiments, participants first completed a passive, sequential reward task that included several high RPEs indicating changes in the underlying distribution of rewards. We then investigated the degree to which high RPEs affected the temporal organization of memories through recognition priming as well as sequence and distance memory measures. We reasoned that if high-RPE events are more strongly bound to the context they were encoded in, then events around the high RPE would be more accessible to one another, resulting in improved priming and better sequence memory. On the other hand, if high-RPE events create new contexts in memory, then events that occurred on either side of a high RPE would be less accessible to one another, leading to less effective priming and sequence memory relative to other pairs of events at the same presentation distance.

We further asked, if high RPEs do create event boundaries, where does this boundary occur? In other words, is the high-RPE event the last of the old context or the first of the new one? The latent cause model would predict that, because the RPE event is predictive of the rewards to follow, it should be the first event of a new context (Gershman et al., 2014). However, recent work suggests that event boundaries lead to the neural reinstatement of events that preceded the boundaries (Baldassano et al., 2017; Ben-Yakov & Dudai, 2011; Ben-Yakov, Eshel, & Dudai, 2013; Sols, DuBrow, Davachi, & Fuentemilla, 2017), which could bind the RPE event to its predecessors. Here, we tested for each of these possibilities by characterizing the associative links between a high-RPE event and its predecessor, as well as those between its predecessor and its successor (i.e., the events around a high RPE, excluding the event itself), in comparison to sequential events that did not cross a high RPE.

To answer these questions, we compared associative and temporal memory for high and low-RPE events using recognition priming (Exp. 1-3) and sequence and distance memory tasks (Exp. 4). We additionally developed a computational model (a variant of the Context Maintenance and Retrieval model; Polyn, Norman, & Kahana, 2009), where high RPEs induce mnemonic separation between rewarding events, and used this model to simulate performance on our experiments and test whether it captured our main behavioral results.

## 2. Overview of experiments

### 2.1 Recognition priming

In Experiments 1-3, we used a recognition priming task to probe whether RPEs influence the degree to which two sequential events are bound in memory. In recognition priming, recognition for an event is better and faster if it is preceded by the event that occurred before it during encoding (Schwartz, Howard, Jing, & Kahana, 2005; Zwaan, 1996). The idea is that retrieval of an item also reactivates items that were associated with it during encoding, either directly, or indirectly via context, facilitating subsequent recognition of those items. This is strongest for the forward sequence (i.e., each cue will reactivate the subsequent one; Howard & Kahana, 2002). Given this, we reasoned that if a high RPE creates an event boundary that separates the high-RPE event from its predecessor, high-RPE events would become less accessible when primed during retrieval, demonstrating less recognition priming. If, instead, high-RPE events are more strongly bound to the previous event, we would expect the RPE event to be more accessible when primed by the preceding event, leading to enhanced recognition priming.

### 2.2 Sequence and distance memory

In Experiment 4 (and its replication), we further tested whether high-RPE events disrupt the integration of sequential events by probing the temporal ordering and perceived distance between them. Contextual changes (both external and internal to an observer) are thought to increase change in one’s internal context, leading to greater perceived time between events (Sahakyan & Smith, 2014). Performance on these measures of temporal memory is modulated by representations in the hippocampus, thought to support the temporal structuring of events in memory (Davachi & Dubrow, 2015): Previous studies have found that greater hippocampal dissimilarity between two events across an event boundary predicts worse sequence memory and larger subjective distances between them (DuBrow & Davachi, 2014; DuBrow & Davachi, 2016; Ezzyat & Davachi, 2014). For sequence memory, we asked participants to indicate which of two items came first, and for distance memory, we asked participants to indicate how far apart the events had been during encoding. If a high RPE signals an event boundary, we would expect worse sequence memory and greater estimated distances for pairs that include or are interrupted by a high-RPE event. On the other hand, if high-RPE events are more bound to the events around them, thereby activating and compressing the sequence of events in memory, we could expect better sequence memory and shorter estimated distances.

## 3. Experiment 1

### 3.1. Method

#### 3.1.1. Participants

Participants were recruited from Amazon’s Mechanical Turk (MTurk), and 35 participants initiated the task (age: 27-67, median = 34; 15 female, 20 male). We first obtained informed consent online, and prior to accessing the task, participants had to correctly answer questions that checked for their understanding of the instructions. All procedures were approved by Princeton University’s Institutional Review Board. We excluded participants if they (a) missed more than 20 memory trials, or (b) had a memory score of less than 0.5 (memory score was determined by A’; Pollack & Norman, 1964). Using these criteria, we excluded 8 participants, which led to a sample of 27 participants.

#### 3.1.2. Task design

Participants completed 6 blocks, each consisting of learning (36 trials in each block), choice (4 trials in each block), and recognition memory phases (42 trials in each block). In the instructions, participants were told they would be exploring six different “rooms” (i.e. blocks), defined by distinct color backgrounds, where they would “find” different photographs and earn 10% of the reward value associated with each photograph. We used a Pavlovian (passive) learning design in order to isolate the effects of changes in reward alone, unconfounded by shifts in responding. In the learning phase, participants passively viewed a sequence of trial-unique images of scenes that were associated with different reward values (Figure 1A). On each trial, participants saw the scene image for 1 second, then were shown the image with its associated value for 2 seconds. The individual values of the scenes fluctuated around a fixed mean (means ranged from 10¢ to 90¢ in steps of 10¢). Participants were encouraged to remember the individual values of the photographs as they would be choosing between them later (after each room), and earning the reward value of the chosen image.

**Figure 1.**
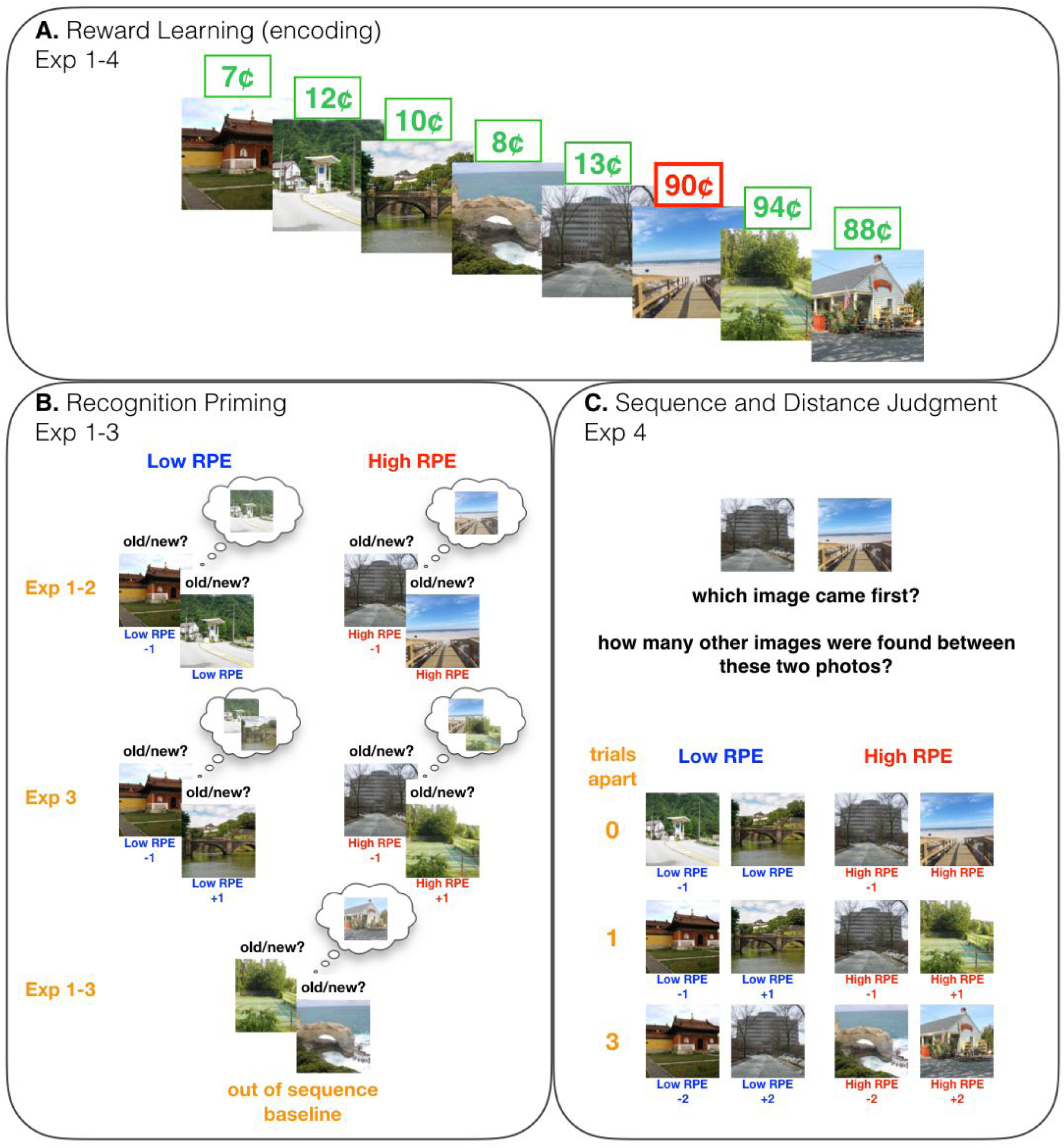
Experimental paradigm. **A**. In all experiments, in each of six blocks, participants first completed a passive reward learning task (the encoding task) where sequences of scenes, each with an associated reward value, were presented. The reward values of the images were contingent on the mean value of the *reward state*, which shifted 4-5 times each block. For illustration purposes, the shift is marked in red, however, it was not distinguished in any way in the actual experiment. **B**. In Exp. 1-3, after reward learning, participants completed a recognition test where they indicated whether a scene was “old” or “new”. We tested for recognition priming of high and low-RPE events, relying on a mechanism by which recognition of an old item (the prime), either directly or indirectly, activates the items that had followed it during encoding (the target), leading to better and faster recognition of target items. Most of the old scenes were presented in pairs that belonged to three different conditions (example stimuli refer to the reward sequence in A): (1) “low RPE”: a pair that was studied consecutively; both items belonged to the same reward state, (2) “high RPE”: a pair that was studied consecutively, however the items belonged to different reward states, (3) “out of sequence” (baseline): the second item in the test pair actually preceded the first item during encoding (i.e., out of order); the items belonged to different reward states. Recognition priming for low and high-RPE pairs was compared to the out-of sequence pairs. In Exp. 1-2, the low and high-RPE pairs comprised items that were directly one after the other during encoding, whereas in Exp. 3, the pairs were separated by another scene during encoding (“+1”), and so the high-RPE (+1) pair did not include the high-RPE event itself. **C**. In Exp. 4 (and its replication), after reward learning, we tested for the temporal memory of two scenes that either belonged to the same reward state (low RPE) or a different reward state (high RPE), and were either 0 (back-to-back), 1 or 3 trials apart. We first asked participants to indicate which of two images came first during encoding (sequence memory), and then for the number of images that occurred between them (distance judgment, scale 0-5). Example pairs (bottom) refer to the reward sequence in A, although unlike the pairs of stimuli presented here, no scene was repeated during testing.

In each room, the mean value of the photographs shifted either four or five times. Participants were told that a shift in the mean value of the photographs indicated they had found a new “collection” of photographs that were more or less valuable than their previous collection. Critically, as a result of these *reward shifts*, participants experienced high positive or negative reward prediction errors whose magnitude ranged from 20¢ to 80¢ (and every 10¢ increment in between; these magnitudes reflect a one-trial difference between current and previous reward). Each participant experienced each magnitude of prediction error 1-2 times, and the number of positive and negative reward shifts was balanced (13 positive and 13 negative high-RPE events across the entire experiment). Within each latent *reward state*, participants experienced at least 5 and at most 9 trials (average = 6.75 trials) where the individual values of the scene images fluctuated around the same mean value (individual reward values never deviated more than 5¢ from the mean value). After learning, within each block, participants completed 4 choice trials that were intended to ensure they paid attention to the values in the passively viewed sequence. On each choice trial, two previously-seen images were presented and the participant chose one, anticipating that the reward value of that image would be added to their payment for participating in the experiment. The 8 images used in the choice test were not used in any other memory test in that block.

#### 3.1.3. Recognition priming

Following the choice test, we tested for recognition priming of pairs that had either been experienced sequentially during encoding or not (Figure 1B). On each recognition trial, participants were asked to indicate “old” or “new” for the presented image (by pressing ‘o’ or ‘n’ respectively). Participants were told to indicate their recognition judgement as quickly as possible. Importantly, the image stayed on screen for 3 seconds regardless of the response time, ensuring that each prime was experienced for the same amount of time.

Recognition trials were comprised of (1) an old scene image, followed by either (a) an old scene image that had followed the prime during learning and belonged to the same reward state (“low RPE”; 4 “priming pairs” within each recognition block, 24 pairs in total) or (b) an old scene image that had followed the prime during learning but belonged to a different reward state (“high RPE”; 4 priming pairs within each recognition block, 24 pairs in total); (2) an old scene item followed by a different old scene image than the one that had followed the prime (“out of sequence”; 4 pairs within each recognition block, 24 pairs total); (3) “single” old scene images, so that participants didn’t always expect two back-to-back old items (“single”; 4 images within each recognition block; 24 images total); (4) new scene images, representing one-third of the images seen during recognition (“new”; 14 images within each recognition block; 84 images total). For the paired test conditions (conditions 1 and 2 listed above) we will refer to the first tested item as the prime and the second tested item as the target. Note also that this pair structure was not disclosed to participants (all test items were presented together in a single sequence).

Recognition priming is evidenced by better memory and faster reaction times in recognizing a target item after correctly retrieving the prime, compared to when the target was preceded by an old item that had not preceded it during encoding. In other words, the baseline for assessing recognition priming in high- and low-RPE pairs was memory accuracy and latency for the out-of-sequence pairs. Although we tested for differences in both hit-rate and response latencies for target items, recognition priming is more consistently observed in response latencies rather than hit-rates (DuBrow & Davachi, 2014; Zwaan, 1996), and so we focused on characterizing (and modeling) recognition latencies within the above four conditions. We were primarily interested in whether recognition priming was enhanced or interrupted for events that had been associated with a high RPE. If a high-RPE event is bound to the event that occurred immediately before it, we would expect faster recognition of the target. On the other hand, if high-RPE events create a boundary in memory during the event itself, we would expect similar reaction times in recognizing the high-RPE target and the out-of-sequence target.

#### 3.1.4. Statistical analysis

All statistical comparisons were conducted using linear or generalized linear mixed-effects models (using lme4 package in R; Bates et al., 2015), treating participant as a random effect for both the intercept and the slope of the tested fixed effect. To test for differences in memory (i.e., hit-rate) between the primed pairs, we analyzed trials where the prime had been correctly remembered; we did this because of prior research indicating that recognition priming only occurs when the prime is itself remembered (Schwartz et al., 2005). When testing for recognition priming in reaction time, we analyzed trials where both the prime and the target were correctly remembered. Reaction times were log-transformed and z-scored within participant.

### 3.2. Results

#### 3.2.1. Recognition memory

We found that the primed targets were better remembered than the out-of-sequence targets, regardless of the RPE condition (*B* = 0.35, *z* = 2.91, *p* = 0.004; low RPE: *B* = 0.38, *z* = 2.75, *p* = 0.006; high RPE: *B* = 0.32, *z* = 2.18, *p* = 0.03; Figure 2A). We did not find a difference in memory between the primed high-RPE and low-RPE images (*B* = −0.04, *z* = −0.30, *p* = 0.76) nor between images in the two non-primed conditions (i.e. the out-of-sequence vs. the “single low RPE” items: *B* = −0.01, *z* = −0.05, *p* = 0.96).

**Figure 2.**
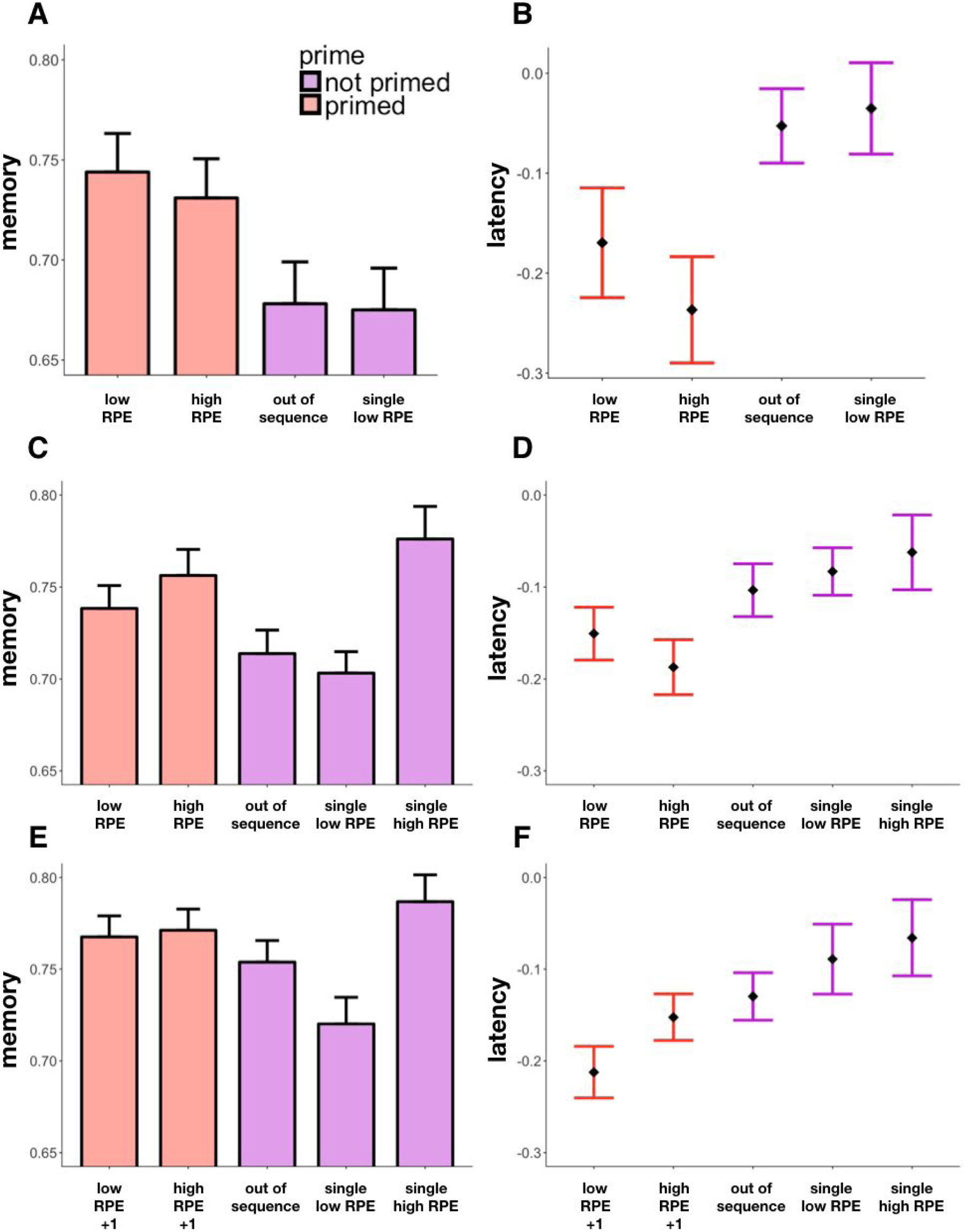
Recognition priming results. For paired targets (“low RPE”, “high RPE” and “out of sequence”), memory is conditioned on correct recognition of the first item in the pair, and response latency is additionally conditioned on correct recognition of the target (i.e., latency is only for “hits” in all conditions). **A**. Exp. 1: Recognition memory as a function of item condition. Memory for the sequentially primed targets (low and high RPE) was better than the out-of-sequence and (unpaired) “single low RPE” targets. **B**. Exp. 1: Response latencies for correct recognition as a function of item condition. Sequentially primed targets were retrieved faster than items that were not sequentially primed. **C**. Exp. 2: Recognition memory as a function of item condition. Memory for the primed high-RPE target was no different than the “single high RPE” target that had not been primed. Thus, memory accuracy did not provide evidence for recognition priming of high-RPE events. **D**. Exp. 2: Response latencies for correct recognition as a function of item condition. Primed high-RPE targets were retrieved faster than the non-primed high-RPE targets and out-of-sequence targets, thereby demonstrating recognition priming for high-RPE events. **E**. Exp. 3: Recognition memory as a function of item condition. Primed targets (where the prime was the item presented two trials before the target during encoding) were not remembered better than the out-of-sequence targets. **F**. Exp. 3: Response latencies for correct recognition as a function of item condition. The high-RPE (+1) target was no longer retrieved faster than the out-of-sequence target, whereas the low-RPE (+1) target was still retrieved faster, demonstrating intact recognition priming. Moreover, latencies for the high-RPE (+1) target were significantly slower than the low-RPE (+1) target. Error bars represent standard error of the mean (SEM).

#### 3.2.2. Reaction time

The primed targets were more quickly recognized than the out-of-sequence targets (*B* = −0.15, *t* = −2.82, *p* = 0.005; Figure 2B), providing additional evidence of recognition priming. This was significant for high-RPE targets (*B* = −0.19, *t* = −3.11, *p* = 0.002), and trending for low-RPE targets (*B* = −0.11, *t* = −1.83, *p* = 0.06). Latencies were moreover no different between the two primed conditions (*B* = −0.08, *t* = −1.16, *p* = 0.25). Additionally, the latencies for correctly recognizing the non-primed targets were not significantly different across conditions (*B* = 0.02, *t* = 0.31, *p* = 0.76).

### 3.3. Discussion

We found better and faster recognition of items that had been primed, including items that were associated with a high RPE. These results suggested that a high-RPE event is bound to its predecessor. However, given that high-RPE items are generally better remembered (Rouhani et al., 2018), it is possible that the generally stronger memory trace is driving the recognition memory results, and not a stronger association with the previous item. We therefore tested in Experiment 2 whether there are differences in the recognition of primed versus non-primed high-RPE items. Specifically, if a high-RPE event is more bound to the preceding event in memory, then we would expect better memory and faster latencies for high-RPE items that are primed versus those that are not primed.

## 4. Experiment 2

### 4.1 Method

#### 4.1.1. Participants

One-hundred participants from MTurk (age: 22-71, median = 35; 46 female, 54 male) were recruited on MTurk. Following the same exclusion criteria stated in Experiment 1, we excluded 17 participants, leaving a final sample of 83 participants.

#### 4.1.2. Task design

Experiment 2 was identical to Experiment 1 except that during the recognition test we additionally included “single” (i.e., not primed) scene images associated with high RPEs. We did this to determine whether high-RPE events lead to better and faster recognition because they are more strongly bound to the previous item (and thus show more recognition priming) or because they are more strongly encoded (i.e., a recognition effect not affected by priming). This led to one fewer high-RPE pair within each recognition block, and 41 trials within each recognition block. Across the experiment, for each participant we tested 18 high-RPE priming pairs, 24 low-RPE priming pairs, 24 out-of-sequence pairs, 16 low-RPE single images, and 8 high-RPE single images.

### 4.2. Results

#### 4.2.1. Recognition memory

We again found that the high-RPE primed items were better remembered than the out-of-sequence items (*B* = 0.24, z = 2.34, *p* = 0.02; Figure 2C); however, we did not find them to be better remembered than high-RPE images that were not primed (*B* = 0.05, z = 0.37, *p* = 0.71). Therefore, we could not conclude that better recognition memory for the high-RPE images was necessarily a result of recognition priming. Additionally, and consistent with previous research (Rouhani et al, 2018), we found the high-RPE (single) items were better remembered than the low-RPE (single) items (*B* = 0.39, z = 3.40, *p* < 0.001).

#### 4.2.2. Reaction time

We replicated our previous observation of faster reaction times in recognizing the primed high-RPE items than the out-of-sequence ones (*B* = −0.08, *t* = −2.11, *p* = 0.03; Figure 2D). Importantly, primed high-RPE images were also recognized more rapidly than the non-primed (single) high-RPE images (*B* = −0.13, *t* = −2.84, *p* = 0.005). The faster recognition of the primed high-RPE items thus reflected intact recognition priming. When testing for a difference between high and low-RPE targets that were primed versus those that were not primed, we did not find an interaction (*B* = −0.05, *t* = −0.76, *p* = 0.45). This indicates that the difference in reaction times for recognizing high-RPE items versus low-RPE items was not different depending on whether those RPE events were or were not primed.

### 4.3. Discussion

We found that high-RPE items were both better remembered overall, and were also primed (at least with regard to reaction time) by recognition cues. From this, we concluded that high-RPE items were, in fact, linked with the items that had occurred before them during encoding, providing no evidence of an event boundary for that association. However, it remained possible that the boundary occurs
across rather than during the high-RPE event. To investigate this possibility, we next tested for priming between pairs that had one item in between them during encoding. This allowed us to exclude the high-RPE item itself and determine whether we see diminished priming for events across a high-RPE versus those across a low-RPE event.

## 5. Experiment 3

### 5.1. Method

#### 5.1.1. Participants

We again recruited 100 participants on MTurk (age: 20-66, median = 33.5; 39 female, 61 male), and following the exclusion criteria stated in Experiment 1, we excluded 15 participants, leading to a final sample of 85 participants.

#### 5.1.2. Task design

The task structure was the same as in Experiments 1 & 2. During recognition, however, instead of testing pairs that had been presented directly one after the other during learning, we tested recognition priming for pairs that had one item in between them during learning. In other words, the high-RPE priming pair never included the high-RPE event itself, allowing us to test whether the events around a high RPE provide evidence of an event boundary. As before, the image immediately preceding the high-RPE event was the prime, but the target was now the image after the high-RPE image (“high RPE +1”). The low-RPE priming pairs had also been one trial apart during learning (“low-RPE +1”), and were selected from the same reward state. All primed targets were therefore associated with low RPEs. We tested 24 high-RPE priming pairs, 24 low-RPE priming pairs, 24 out-of-sequence pairs, 12 single low-RPE images, and 12 single high-RPE images along with 84 new images, across all 6 blocks of the experiment (42 trials within each recognition block).

### 5.2. Results

#### 5.2.1. Recognition memory

Memory was not significantly better for the primed targets in comparison to the out-of-sequence items (*B* = 0.13, *z* = 1.57, *p* = 0.12; Figure 2E). We again found better memory for high RPE (single) items relative to low-RPE (single) items (*B* = 0.41, *z* = 3.44, *p* < 0.001).

#### 5.2.2. Reaction time

When excluding the high-RPE item itself, we no longer observed a recognition priming effect for pairs that spanned a high-RPE event (compared to out-of-sequence, *B* = −0.02, *t* = −0.53, *p* = 0.60; Figure 2F). We nevertheless did see recognition priming for pairs that spanned a low-RPE event (compared to out-of-sequence, *B* = −0.08, *t* = −2.42, *p* = 0.02). Moreover, there was now a difference between the latencies of the high- and low-RPE pairs where the high RPE +1 items were more slowly recognized than the low-RPE +1 targets (*B* = 0.06, *t* = 1.89, *p* = 0.05).

### 5.3. Discussion

The results of Experiment 3 provide evidence that high RPEs serve as an event boundary. The slower latencies in recognizing the item that followed the high-RPE prime, which were now similar to the out-of-sequence pairs and significantly slower than the low-RPE pairs, indicated decreased recognition priming. With this initial evidence of an event boundary, we next tested whether events around a high RPE demonstrate other behavioral markers of event boundaries. For this, we asked whether high-RPE events disrupt the temporal organization of events in memory, leading to worse sequence memory and larger perceived distances between item-pairs that include a high-RPE event versus those that do not.

## 6. Experiment 4

### 6.1. Method

#### 6.1.1. Participants

For the first set of this experiment, we recruited 50 participants on MTurk (age: 24-61, median = 38; 26 female, 24 male). We excluded participants if they missed more than 15 trials, which led to the exclusion of 1 participant and a final sample of 49 participants.

Subsequently, we ran an additional sample of 80 participants as a pre-registered replication of this experiment (for pre-registration, see Rouhani, 2018). The replication sample size was chosen on the basis of a simulation-based power analysis of the effect seen in the initial sample, which indicated we would have sufficient power (80% probability) of replicating the results with 50 participants. Following common practice of testing around 1.5x the indicated sample size for replication studies, we thus recruited 80 participants on MTurk (age: 24-68, median = 38, 38 female, 42 male), and excluded 3 participants who missed more than 15 trials, leaving a final sample of 77 participants.

#### 6.1.2. Task design

The task structure was the same as in Experiments 1-3; however, instead of testing for recognition memory, here we tested participants’ sequence memory and distance judgements for images seen during learning. Worse sequence memory and larger estimated distance between items are considered as evidence of an event boundary in memory (Davachi & DuBrow, 2015). We instructed participants to pay attention to the sequence of images during learning as they would later be asked to order them. After the learning and choice sections in each block, participants were presented with two old scene images on the screen (left/right order counterbalanced), and were asked to indicate which image came first (“sequence memory”) and then to estimate how many other images were found between the two (from 0-5; “distance judgment”; Figure 1C). Within each block, participants completed 12 sequence and distance judgment trials. The two scene images either spanned (or even included) a high-RPE event (“high RPE”; 48 total), or were from the same reward state (“low RPE”: 48 total). Additionally, the high/low-RPE manipulation was crossed with a distance manipulation: the pairs had either been presented directly one after the other (“0 between”: 24 total), had one item in between them (“1 between”: 24 total), or had three items in between them (“3 between”: 24 total) during learning. Note that the “0 between” high-RPE pairs included the high RPE event and the event that immediately preceded it. The “1 between” high-RPE pairs included the events immediately preceding and following a high-RPE event, and the “3 between” high-RPE pairs included the second event before and the second event after the high-RPE event.

### 6.2. Results

#### 6.2.1. Sequence memory

We found better sequence memory for pairs within the same reward state than across a high RPE (*B* = 0.25, *z* = 3.46, *p* = 0.0005; Figure 3A), and replicated this main effect in the second sample (*B* = 0.17, *z* = 2.97, *p* = 0.003; Figure 3B). Interestingly, when the high RPE was included in the pair (“0” trials apart), there was no difference in sequence memory between the pair types (first set: *B* = −0.03, *z* = −0.26, *p* = 0.79; replication set: *B* = −0.05, *z* = −0.52, *p* = 0.61). The difference in sequence memory was instead carried by pairs that did not include the high-RPE event itself, i.e. the pairs that had 1 item in between them (first set: *B* = −0.35, *z* = −2.79, *p* = 0.005; replication set: *B* = −0.19, *z* = −1.87, *p* = 0.06), and 3 items between them (first set: *B* = −0.36, *z* = −2.95, *p* = 0.003; replication set: *B* = −0.27, *z* = −2.88, *p* = 0.004).

**Figure 3.**
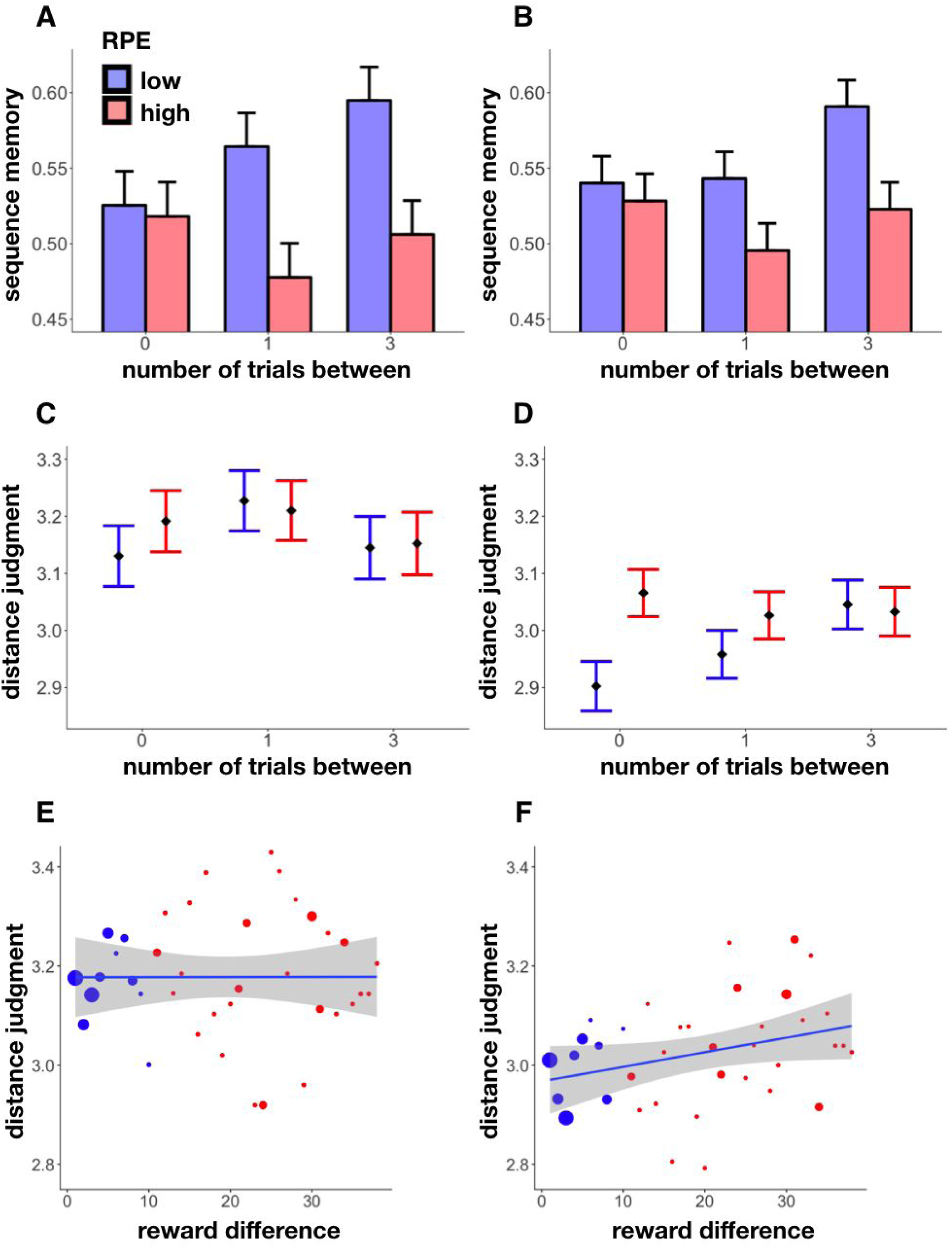
Sequence and distance memory results. **A-B**. Sequence memory in Exp. 4 (A) and its replication (B) as a function of RPE event and presentation distance (number of trials) within scene pairs. Sequence memory for pairs that spanned a high-RPE event was worse; this was driven by pairs that did not include the high-RPE event itself (i.e., pairs that were 1 or 3 trials apart). **C-D**. Distance judgement in Exp. 4 (C) and its replication (D) as a function of RPE event and presentation distance (number of trials) within scene pairs. High-RPE events were perceived as more distant from each other only in the replication experiment, a result driven by pairs that included the high-RPE event itself (i.e., 0 trials apart). **E-F**. Distance judgement as a function of the reward difference between scenes in Exp. 4 (E) and its replication (F). In the replication experiment, we found that greater reward difference between scenes, which was a proxy for the magnitude of the RPE event that had occurred between them, led to greater perceived distance. Size of the dots reflects the size of that sample. Shaded regions reflect 95% confidence intervals. Error bars represent SEM.

#### 6.2.2. Distance memory

We did not find that high RPEs influenced distance judgments in the first dataset (*B* = −0.01, *t* = −0.52, *p* = 0.61; Figure 3C). To further assess whether the magnitude of the RPE influenced perceived distance, we correlated distance judgments with the reward difference between the pair of items within a pair (which is a proxy for the magnitude of any intervening RPE event, since item values were roughly stable on each side of a high-RPE event). We did not find this measure to predict perceived distance either (*B* = 0.03, *t* = 0.89, *p* = 0.38; Figure 3E).

In the larger replication dataset, however, we did find two main effects and an interaction between RPE event and presentation distance in modulating distance judgments (Figure 3D). Here, perceived distance was higher when the pair included/spanned a high (vs. low) RPE event, and it was also higher when there were more trials between two images in a pair (RPE: *B* = 0.11, *t* = 3.12, *p* = 0.002; trials apart: *B* = 0.03, *t* = 2.43, *p* = 0.02). These two effects interacted such that the high-RPE effect was strongest for items that were closer together (*B* = −0.04, *t* = −2.05, *p* = 0.04), and in particular for the pairs that had included the high-RPE item itself (0 trials apart; *B* = 0.12, *t* = 2.91, *p* = 0.004). We also found that the greater the reward difference between the two images, the greater the perceived distance (*B* = 0.04, *t* = 2.83, *p* = 0.005; Figure 3E). This effect was largely driven by the pair that included the high RPE (0 trials apart: *B* = 0.07, *t* = 2.45, *p* = 0.01; 1 trial apart: *B* = 0.05, *t* = 1.85, *p* = 0.06; 3 trials apart: *B* = 0.01, *t* = 0.56, *p* = 0.58).

### 6.3. Discussion

In Experiment 4 and its replication, we again found that high-RPE events act as event boundaries by interrupting the sequential integration of events into memory, leading to worse sequence memory for events around the high RPE. Interestingly, there were no differences in sequence memory for the pair that included the high-RPE item itself, again suggesting that the high-RPE event is associated with its predecessor (similar to intact recognition priming for high-RPE events in Experiments 1-2). We only found an effect of high RPEs on perceived distance in the replication dataset: High RPEs led to greater perceived distance, and (relatedly) greater differences in reward value between the two items were associated with greater perceived distance; importantly, these effects were most reliably present for the “0 between” condition, where the pair included the high-RPE event. Given that the effects of RPE on sequence memory showed the opposite pattern (i.e., effects on sequence memory were largest for the pairs that did not include the high-RPE event, and absent for the pairs that did include the high-RPE event). This qualitative difference suggests a potential dissociation between the mechanisms supporting sequence and distance judgments. Nevertheless, as we did not find these distance effects in the first dataset, they require further investigation and replication.

## 7. Computational model

### 7.1. Overview

To explore potential mechanisms for our findings, we developed a variant of the Context Maintenance and Retrieval model (CMR; Polyn et al., 2009; for other variants, see CMR2: Lohnas, Polyn, & Kahana, 2015; eCMR: Talmi, Lohnas, & Daw, 2019), and tested whether our behavioral results can be explained by a model in which high RPEs induce mnemonic separation between events. In our model, experienced events are temporally linked through a slowly drifting internal “context”, where features of the experienced items update the context representation (Howard & Kahana, 2002). We posit that high RPEs temporarily increase the context drift rate (i.e., the extent to which the high-RPE event updates context), thereby creating a large shift between the context representation of events experienced prior to the high RPE and those experienced after it. We show that this discontinuity can explain our findings of reduced recognition priming (Exp. 3) and worse sequence memory (Exp. 4) after high RPEs.

We used the model to simulate both recognition priming and sequence memory. To simulate recognition priming, we first presented a recognition prime to the model, which triggered an update to the model’s context representation. Next, the recognition target was presented to the model. Importantly, activation was allowed to spread back from the context representation (which had been updated by the prime) to the representation of the target; this spreading activation affected the latency with which the target was recognized (for details, see *7.6* below). To simulate sequence memory, we calculated the similarity of the context representations that were linked to the to-be-judged items, and used these similarities to predict participants’ sequence memory scores in the different conditions (for details, see *7.7* below).

### 7.2. Representational structure

The model includes two layers, a feature layer (*F*) and an internal, temporal context layer (*C*), both of which contain the same number of units. External events (happening at time *i*) activate a single localist feature in *F* (**f**_*i*_), and these activations spread up from *F* to *C* (the context layer at time *i* is denoted as **c**_*i*_) via a feature-to-context matrix (*M*^*FC*^) that updates context during both the initial encoding phase and the test phase. During retrieval, activations spread back down from *C* to *F* via a context-to-feature matrix (*M*^*CF*^) that guides memory search (Figure 4). We represent different events as orthogonal unit vectors (“one-hot”). Although the CMR uses an additional “source layer” to tag explicit contextual shifts (such as different encoding tasks), in our model we did not use this layer to tag different reward states. This is because changes in the reward distribution were latent to the participant (and thus also to the model).

**Figure 4.**
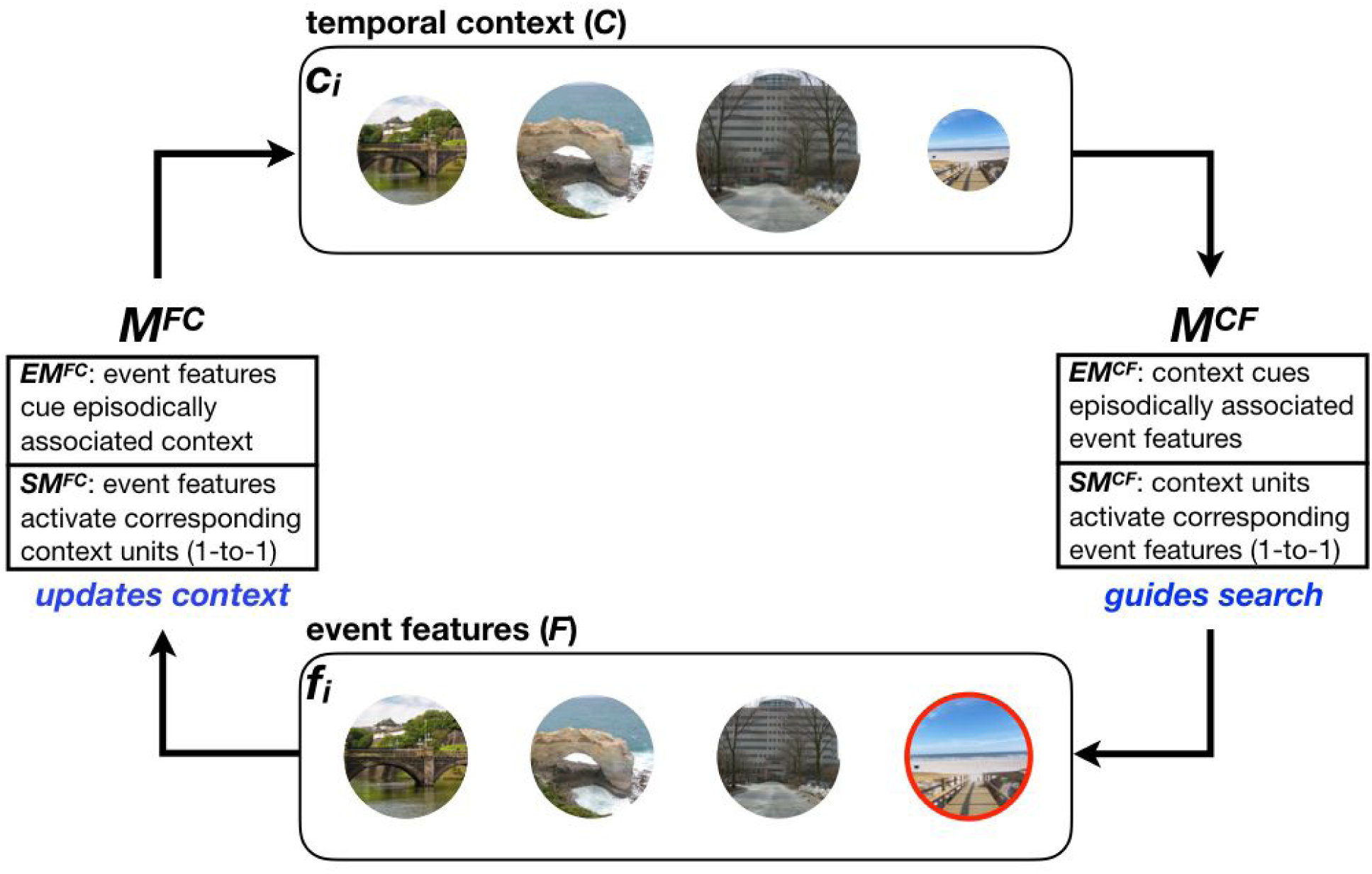
Model structure. The model has two layers: a feature layer (*F*) and a temporal context layer (*C*) that interact through two associative matrices: a feature-to-context matrix (*M*^*FC*^) that updates context and a context-to-feature matrix (*M*^*CF*^) that guides search. Each matrix is a composite of an episodic (*EM*^*CF*^, *EM*^*FC*^) and semantic matrix (*SM*^*CF*^, *SM*^*FC*^). The episodic matrices represent the episodic associations formed between *F* and *C* during encoding, whereas the semantic matrices contain one-to-one connections between features in *F* and the corresponding units in *C*. When an event is “experienced” (during encoding) or “remembered” (during retrieval), its corresponding unit *f*_*i*_ is activated in *F*, and activation spreads up to *C* via *M*^*FC*^. Specifically, *EM*^*FC*^ updates *C* with contexts that were previously (episodically) linked to *f*_*i*_ (“mental time travel”), and *SM*^*FC*^ updates *C* by activating the unit in the context layer that directly corresponds to *f*_*i*_ (e.g., if *f*_*i*_ is the third unit in the feature layer, *SM*^*FC*^ activates the third unit in the context layer). During retrieval, activation spreads down from *C* to *F* via *M*^*CF*^. Specifically, *EM*^*CF*^ activates units in *F* that were previously (episodically) linked to contexts that match the current state of *C* (“episodic retrieval”), and *SM*^*CF*^ activates units in *F* proportionally to how active the corresponding units are in *C* (“direct readout”). Units in *F* then compete for retrieval. The figure depicts the state of the model at time point *i* = 4: The first three items (from left to right) were presented successively on previous trials, and are therefore active in context (more recently experienced items are more active in *C*, as reflected here by the size of the circles); the fourth item (outlined in red) is being presented in the feature layer. This feature-layer representation of the fourth item will be episodically associated with the context shown here; on the *next* time step it will be used to update the state of *C* (via *M*^*FC*^) and the cycle will begin again.

Each associative matrix was made up of an episodic and a semantic component, meaning that *M*^*CF*^ comprised a weighted average of episodic (*EM*^*CF*^) and semantic (*SM*^*CF*^) weight matrices, and likewise *M*^*FC*^ comprised a weighted average of *EM*^*FC*^ and *SM*^*FC*^ (we modeled the weights of each matrix separately, see *7.5-7.6* below). As in TCM and CMR, the episodic matrices are updated during encoding to store associations between active feature representations in *F* and context representations in *C*. The semantic matrices contain one-to-one connections between a unit in *F* to its corresponding unit in *C* (concretely, they are identity matrices).

### 7.3. Updating temporal context and associative matrices during reward learning

Prior to the reward learning phase, *C* and the episodic associative matrices (*EM*^*CF*^ and *EM*^*FC*^) are initialized to 0. When an item is activated in *F* during the reward learning phase, the activation spreads up from *F* to *C* via *M*^*FC*^ where the input to *C* is calculated as follows:

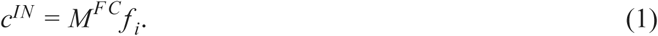

The vector c^IN^ is then normalized to be of unit length, and then context is updated as follows (as in TCM and CMR):

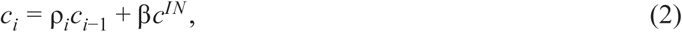

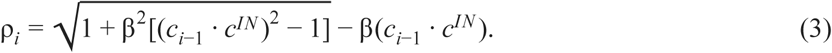

Here, β defines the degree to which the active feature causes the context to “drift” -- the larger the value of β, the more the active feature will be inserted into the context, crowding out other active events in *C*. We allowed for two distinct drift values, β, the standard drift (implemented for low-RPE events), and *d*, a higher level of drift for high-RPE events. This approach (i.e., increased drift in response to high-RPE events) is in line with how contextual disruptions due to salient changes have been previously modeled (Horner et al., 2016; Polyn et al., 2009; Siefke et al., 2019). We moreover use *d* for the first item presented to the network as a way of capturing classic primacy effects in memory (i.e., the higher probability of retrieving the first item in a sequence; see *7.5* for further discussion of how primacy is modeled here, compared to how it is usually modeled in CMR).

The two episodic associative matrices are updated through Hebbian outer-product associative learning. α represents the learning rate for that update:

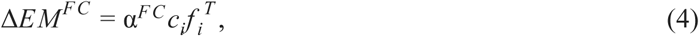

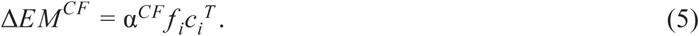

Importantly, in our version of the model, on each time step, the following order-of-operations applies: First, the feature vector is updated based on the current event; next, the episodic matrices are updated; and finally the context vector is updated. The consequence of this order-of-operations is each event is inserted into the *following* event’s episodic context (but not its own episodic context). For example, at the end of the fourth time step, the fourth item will be inserted into the context layer; at the start of time step 5, the fifth item’s feature-layer representation will be activated, at which point it will be episodically associated with the current state of the context layer (where the *fourth* item’s context-layer representation is active). Next, the fifth item’s context-layer representation is activated, and the cycle begins again.

### 7.4. Simulating free recall

Although we did not collect our own free recall data, we calibrated the model by running free-recall simulations, using the following procedure. First, after the learning trials, we simulated the intervening time period before the memory test by presenting 15 randomly-generated “distractor” events. This allowed active features in *C* to substantially drift from the encoding period, thereby capturing the putative drift occurring between the end of the learning phase and the start of the test phase. These distractor events did not compete during retrieval.

The associative matrices at recall were each calculated as a weighted average of their episodic and semantic components:

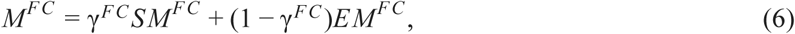

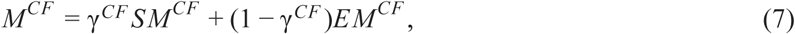

As in CMR and TCM-A, the recall period was governed by a leaky, competitive accumulation process where experienced events accumulated activation until one passed a threshold and “won” the competition. The following calculates the input to the accumulators:

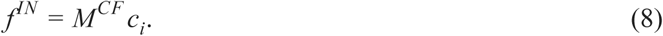

Which then guides the below competition dynamics:

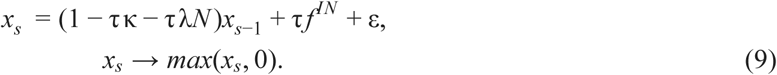

Here, x is a vector with units corresponding to each element in the feature layer (**f**^**IN**^), and *s* indexes the step in the accumulation process (units are initialized to 0, and cannot take on negative values, second line of Eq. 6). The parameters governing the competition are τ, the time constant determining the rate of accumulation, κ, the decay rate for active items, and λ, the lateral inhibition parameter which scales the strength of inhibitory matrix, *N*; ε adds gaussian noise to the decision process (drawn from a random normal distribution with mean zero and standard deviation η). This accumulation process proceeded until one of the elements passed a threshold of 1, at which point the winning item’s feature was reinstated in *F*, and its encoding context was reactivated:

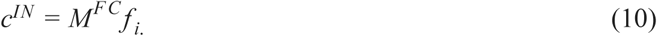

The reactivated context was then used to update the current context vector following Eq. 2. Subsequently, **f**^**IN**^ was updated and the accumulation process restarted with x_(1)_ = 0. Previously retrieved items were allowed to continue competing in the accumulation process, but were prevented from passing the retrieval threshold.

### 7.5. Model calibration

Before simulating our experiments, we determined which parameter values to use by identifying combinations that replicate canonical findings in free recall tasks; namely, the higher probability of recalling the first item (“primacy”) and the last item (“recency”) in a given context, along with contiguity effects (increased likelihood of recalling items that were studied close together in time, with a bias towards forward transitions; Howard & Kahana, 2002). We identified these parameters by feeding our network distinct events (orthogonal one-hot vectors) and running network simulations for all value combinations of the following four parameters (ranging from 0-1, in increments of 0.05; 100 simulations for each combination): (1) *d*, context drift for primacy events (and for high-RPE events, in the recognition simulations presented later); (2) β, context drift for non-primacy events (and for low-RPE events in the recognition simulations); (3) γ^*CF*^, the relative weight assigned to the semantic vs. episodic components in *M*^*CF*^; and (4) γ^*FC*^, the relative weight assigned to the semantic vs. episodic components in *M*^*FC*^. All other parameter values were taken from Polyn et al. (2009; see Supplemental Material). We generated serial position curves and conditional response probability curves for each run, and filtered the parameter values based on whether they generated characteristic features of these recall curves. Specifically, in the serial position curves, the parameter values we chose generated primacy (higher recall of the first item relative to the subsequent one) and recency effects (higher recall of the last item relative to the preceding one). When simulating contiguity effects, we looked for parameter values that resulted in greater sequential recall of events that were neighboring during encoding, with an increased likelihood of forward recall (thereby matching the pattern that is typically observed in free recall; Howard & Kahana, 2002).

We found that recency and contiguity effects were obtained across a fairly wide range of parameters in the model (as has been shown in previous work with TCM and CMR; Howard & Kahana, 2002; Polyn et al., 2009). Primacy effects were obtained across a more narrow range of parameters. Specifically, to obtain primacy effects we needed to have a relatively high drift rate for primacy items (*d*) compared to the drift rate for non-primacy items (β), as well as a strong contribution of the semantic matrix to both *M*^*CF*^ and *M*^*F C*^ (i.e., high values of γ^*CF*^ and γ^*FC*^). This configuration of parameters allowed primacy effects to arise in the following manner: When the primacy item is present, it is strongly inserted into context, due to the high value of context drift (*d*) that we assigned to primacy items, and the high contribution of the semantic matrix to *M*^*F C*^. Because the primacy item is strongly inserted into context, it is still present in context (i.e., its unit’s activation has not fully decayed away) at the time of test. Because of the strong contribution of the semantic matrix to *M*^*F C*^ (which supports “direct readout” of active items in context back into the feature layer), the fact that the primacy item is still active in context leads to increased activation of that item back in the feature layer (via the aforementioned “direct readout” mechanism; see *7.8* for how these matrices interact during our recognition priming simulations). Note that this way of modeling primacy is different from how primacy is handled in CMR -- in Polyn et al. (2009), primacy items are assigned a higher learning rate (for forming episodic context-to-feature associations) but the drift rate is the same for primacy and non-primacy items. A key goal of our modeling exercise was to assess if we could model our own experimental results and also classic recall effects (e.g., primacy) only through drift manipulations and not through learning rate manipulations; we return to this point in *8.2* below.

As a result of these initial simulations, we selected the following parameter values: *d* = 0.8; β = 0.6; γ^*CF*^ = 0.75; γ^*FC*^ = 0.70. We subsequently ran the recognition priming and sequence memory procedure detailed below (see *7.6*-*7.7*) using these parameters. For recognition priming, we ran 10,000 simulations for each condition, and for sequence memory, we ran a single simulation for each condition since dynamics during encoding are deterministic.

### 7.6. Recognition priming

To simulate our recognition priming results, we used the following procedure: After the initial learning phase and presentation of filler items (see *7.3-7.4*), we presented a “recognition prime” to the network by activating the “one-hot” feature vector that represents that event. After the prime’s representation was activated in *F*, activation was allowed to spread up from *F* to *C* via *M*^*FC*^. The *EM*^*FC*^ component of *M*^*FC*^ updates the context vector with the prime’s episodic context (i.e., the context linked to the prime at encoding; this is the process commonly referred to as “mental time travel”, since it makes the context at test resemble the context when the prime was studied; Kragel, Morton, & Polyn, 2015; Tulving, 1984). The *SM*^*FC*^ component of *M*^*FC*^ allows for the prime itself to be inserted into *C* (see *7.8-7.9* for more description on how these matrices interact during retrieval).

Note that prior studies have found that recognition priming is only obtained when the prime is successfully recollected at test (Schwartz et al., 2005). Our allowing activation to spread from the prime’s feature-layer representation to *C* via *EM*^*FC*^ corresponds to an assumption that the prime was (itself) successfully recollected; this assumption is justified because -- in the priming data that we set out to model -- we only analyzed trials where the prime was successfully remembered (so the assumptions of the model match the structure of our analysis; see *3.1.4*).

After context was updated by the prime, the recognition trial was simulated. Here, activation was allowed to spread down from *C* to *F* via *M*^*CF*^. *EM*^*CF*^ modulates item activation as a function of the match between each item’s episodic context and the current context, and *SM*^*CF*^ provides a “direct readout” of activations from *C* to *F* (e.g., if the fifth unit in *C* is active, activity spreads directly down to the fifth unit in *F*). We then allowed the competition dynamics to unfold. To simulate the fact that the recognition target is presented perceptually, we boosted the activation of the target event by in *F* by 0.75 at the start of the competition; this had the effect of ensuring that the target event would be the winner of the competition, but still allowed for variance in recognition latency. We extracted recognition latencies for the target item and compared them with the empirical recognition data.

We tested target items matched to our experimental conditions, and ran simulations for each condition separately. For the simulation of Experiment 2, the “low RPE” target was the low-RPE event that had been studied directly after the prime and the “high RPE” target was the high-RPE event that had been studied directly after the prime; for the simulation of Experiment 3, the low-RPE (+1) target was a low-RPE event that had been studied two events after the prime, and the high-RPE (+1) target was a low-RPE event that had been studied two events after the prime (with the high-RPE event having occurred between the prime and the target). The “out of sequence” target was always an event that had been studied before the prime (3 trials apart). For conditions where there was no prime (“single high-RPE” and “single low RPE”), we did not present a prime to the model prior to simulating target recognition -- in this case, the state of *C* at the start of target recognition only reflected the effects of the reward learning phase and the distractor items (but not the prime); otherwise, the procedure was the same as in primed trials.

### 7.7. Sequence memory

To simulate sequence memory judgments, we compared the similarity of the context representations of the two events being judged, assuming -- in line with previous work -- that greater similarity predicts better sequence judgments (DuBrow & Davachi, 2014; Ezzyat & Davachi, 2014; Horner et al., 2016). Specifically, we extracted the context vectors that had been episodically linked to each of the events that we probed during sequence memory; i.e. pairs that either spanned (or included) a high RPE or not, and were 0, 1 or 3 trials apart during encoding. We computed the Pearson correlation between the context representations of each tested pair. Within each distance condition, we measured the difference in correlation between pairs that involved a low or high RPE, which gave a similarity difference score (between high and low-RPE pairs) within each distance condition. We next used these similarity difference scores to calculate a memory score in the following way: First, we took as baseline the participant’s empirical memory score averaged across conditions. We then took 1/2 of the difference in similarity score as an offset to the baseline memory score. For high-RPE trials, we multiplied the baseline by 1 - the offset, whereas for low-RPE trials, we multiplied the baseline by 1 + the offset, consistent with worse sequence memory for high-RPE pairs and better sequence memory for low-RPE pairs. We then averaged the simulated sequence memory scores across participants and within each condition.

### 7.8. Simulation results

During the initial encoding (i.e., reward learning) phase, our use of a higher drift rate for high-RPE events created a discontinuity in the mental contexts associated with events that occurred before the high RPE event versus those that occurred after it. We tested how this representational “event boundary” affected recognition priming in simulations of Experiments 2 and 3. Experiment 2 (Figure 5A-B) tested pairs of events that were consecutively-encoded during the reward-learning phase -- call these events *n* and *n*+1 (referring to their adjacent positions during learning). For some pairs, event *n*+1 was a high RPE event (“high RPE”), and for other pairs, event *n*+1 was a low RPE event (“low RPE”). As noted in *7.4*, the model is set up such that (during reward learning) each item becomes part of the *next* item’s episodic context (i.e., item *n* is strongly active in the context layer when item *n*+1 is activated in the feature layer; see Figure 4). At test, when item *n* is presented as a prime (by activating its representation in the feature layer), activation spreads up to item *n*’s representation in the context layer (via the influence of *SM*^*FC*^). Next, activation is allowed to spread back down to the feature layer via *M*^*CF*^. Here, the influence of *EM*^*CF*^ is crucial -- the effect of this matrix is that items whose context at study matched the current context are activated in the feature layer. Crucially, because item *n* was part of item *n*+1’s context at study, the effect of *EM*^*CF*^ in this situation is to allow activation to spread from the “item *n*” unit in the context layer to the “item *n*+1” unit in the feature layer. This spreading activation allows the “item *n*+1” unit to cross threshold sooner when item *n*+1 is presented as a recognition target, thereby giving rise to the recognition priming effect.

**Figure 5.**
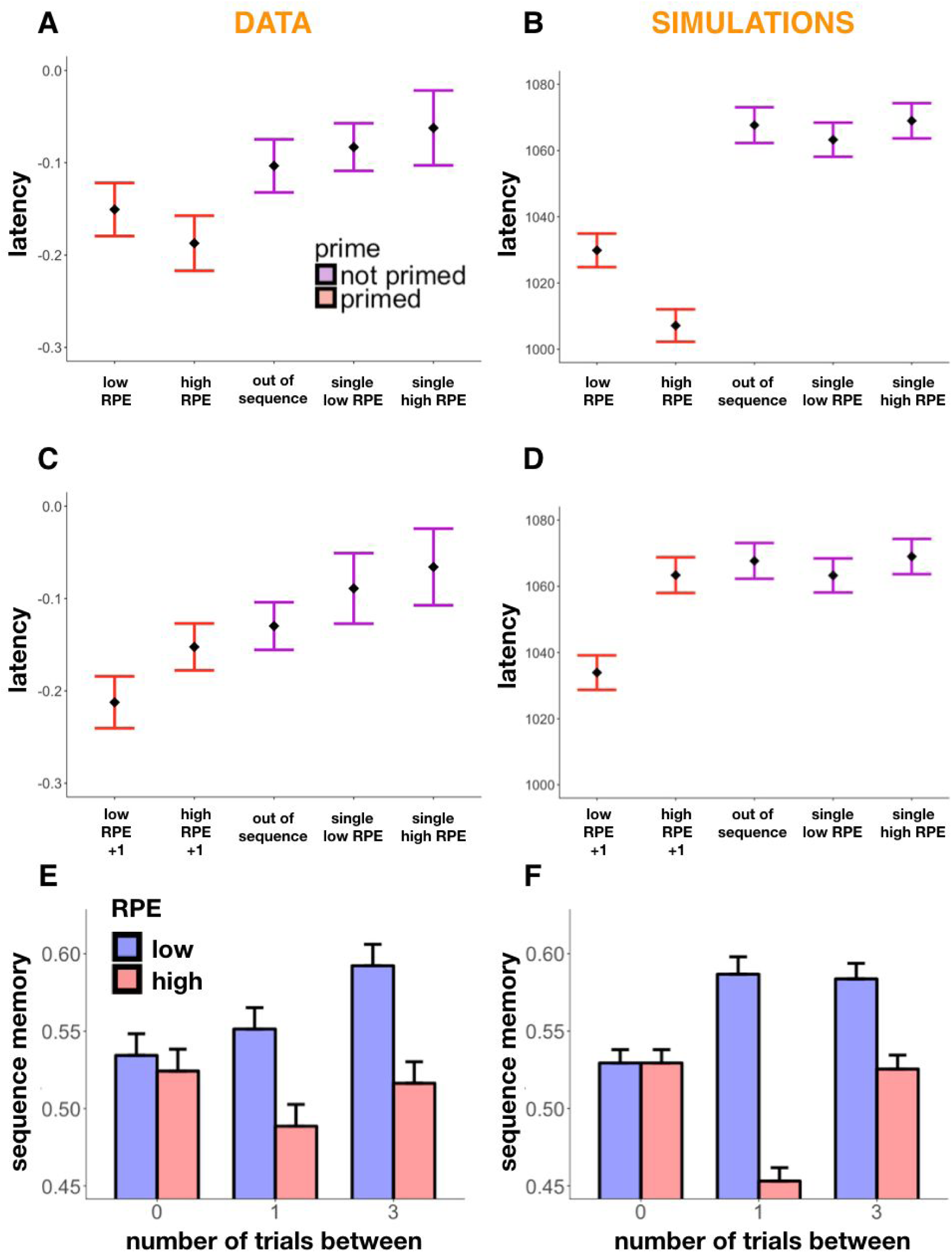
Simulation results along with behavioral results. **A-B**. Recognition latencies as a function of item condition in Exp. 2 (A) compared to model simulations (B). High- and low-RPE targets are retrieved faster than the out-of-sequence targets. In the simulations there is, moreover, an interaction between priming condition and RPE, such that high-RPE targets are retrieved faster than primed low-RPE and this difference depends on whether targets were or were not primed. **C-D**. Recognition latencies as a function of item condition in Exp. 3 (C) compared to model simulations (D). In both the data and the simulations, the high-RPE +1 target no longer shows recognition priming (i.e., it is no longer retrieved faster than the out-of-sequence target) but the low-RPE +1 target shows robust recognition priming. **E-F**. Sequence memory as a function of RPE event and presentation distance (number of trials) within scene pairs, in Exp. 4 and its replication (results averaged across both) (E) and its model simulation (F). Sequence memory for pairs that were not presented adjacently at encoding (i.e., 1-trial-between, 3-trials-between) is worse if the pair spans a high-RPE event; sequence memory for pairs that were presented adjacently at encoding (0-trials-between) is unaffected by whether one of the items is a high-RPE event. Error bars represent SEM.

This priming effect is present in the model for both high-RPE primed targets and low-RPE primed targets, but it is larger in magnitude for high-RPE targets than low-RPE targets (*F*(2,29997) = 35.52, p < 0.001; Tukey-corrected pairwise comparisons: high RPE versus out of sequence, p < 0.001, low RPE versus out of sequence, p < 0.001, high RPE versus low RPE, p = 0.005. Moreover, there was a significant interaction in the retrieval of high and low-RPE targets that were primed versus those that were not, indicating that priming led to the faster retrieval of the high-RPE target relative to the low-RPE target, *F*(1,39996) = 7.75, p = 0.005). The difference in priming effects (in the model) between high-RPE and low-RPE targets is caused by the influence of *SM*^*CF*^ at retrieval. In addition to the effects of *EM*^*CF*^ (described above), *SM*^*CF*^ provides a “direct readout” of which items are active in the context layer. Because of the greater drift associated with high-RPE items, high-RPE (vs. low-RPE) items end up being more strongly active in context (even at the time of test). This extra activation in context translates (via the influence of *SM*^*CF*^) into greater activation of the high-RPE target in the feature layer, which further speeds recognition for high-RPE items, boosting the level of recognition priming.

In Experiment 3 (see Figure 5C-D), primed target items were studied two items after the prime during the learning phase (i.e., with one event in between); sometimes the event interposed between prime and target during learning was a high-RPE event, and sometimes it was a low-RPE event. For the purpose of explaining what happens in the model on these trials, call the prime item *n-1*, the interposed item *n*, and the target item *n*+1. First, consider the condition where the interposed item was a low-RPE event. In this case, during learning, item *n*-1 (the prime) is still strongly active in context when item *n*+1 (the target) is studied, so the prime’s representation in context gets linked to the target’s representation in the feature layer. Because of this link, the usual mechanisms of recognition priming (as described in the preceding paragraph) still apply. Next, consider the condition where the interposed item was a high-RPE event. Because of the higher drift rate for high-RPE items, the effect of (strongly) inserting high-RPE item *n* into context is to “push out” the representation of item *n-1* from the context layer. Because item *n-1* (the prime) is no longer strongly active in context when item *n*+1 (the target) is studied, the crucial episodic link between the prime (in context) and the target (in the feature layer) is not formed, eliminating the recognition priming effect (*F*(2,29997) = 11.85, p < 0.001; Tukey-corrected pairwise comparisons: high RPE versus out of sequence, p = 0.84, low RPE versus out of sequence, p < 0.001, high RPE versus low RPE, p < 0.001). Finally, there was a significant interaction in the retrieval of primed high and low-RPE items between experiments, such that priming of the high-RPE event itself (Exp. 2) is enhanced whereas priming of the high-RPE +1 event (Exp. 3) is interrupted relative to the low-RPE items (*F*(1,39996) = 25.68, p < 0.001).

In the simulation of Experiment 4 (Figure 5E-F), we captured the key characteristics of our sequence memory data. As described above, the strong context drift induced by the high-RPE event at encoding induces a discontinuity in context between items that precede the high-RPE event and items that follow the high-RPE event. Importantly, no such discontinuity in context exists between the items preceding the high-RPE event and the high-RPE event *itself* (i.e., the strong drift induced by the high-RPE event only affects the context for the *following* event; see *7.3* for discussion of this point). The effect of the high-RPE-induced context shift is therefore to reduce contextual similarity when the to-be-judged probes surrounded the high-RPE event during learning (1-trial-apart and 3-trial-apart conditions), resulting in reduced simulated sequence memory performance (1 trial apart: *t*(125) = −52.27, *p* < 0.001, and 3 trials apart: *t*(125) = −58.09, *p* < 0.001). However, since there is no contextual “gap” between the high-RPE event and the preceding event, simulated sequence memory is not negatively affected in this (0-trial-apart) condition.

### 7.9. Discussion of simulation results

Our model, with parameters chosen to generate canonical free recall dynamics, was able to capture the signature effects of our recognition and sequence memory tasks. In our simulation of Experiment 2, we found that feeding the network recognition primes led to the faster retrieval of target items that had come directly after the primes during the initial reward learning phase (i.e., the low- and high-RPE targets) as compared to targets that were out of sequence or were not primed (single items). Recognition priming was especially strong for high-RPE items, whose higher activation in *C* led to faster retrieval times as compared to low-RPE targets. Although we did not observe significantly faster retrieval times for high-RPE versus low-RPE events in Experiments 1 and 2, the simulation results suggest that the numerical difference in their latencies may reflect an actual effect, which may reach significance with sufficient power.

In our simulation of Experiment 3, the prime and the target always had one event (either high-RPE or low-RPE) between them. Our model captured the lack of recognition priming in the high-RPE condition by creating a contextual discontinuity after the high-RPE item, thereby “breaking” the contextual link between the prime and the target. The same mechanism allowed us to simulate the key result from Experiment 4: impaired sequence judgments when the to-be-judged items surrounded a high-RPE event during learning.

In introducing the simulations, we identified four parameters of interest, namely the drift rates for high-RPE and low-RPE events at encoding and the episodic and semantic proportions of the associative matrices. The effects of drift rate on model results ended up being fairly straightforward: *d* (the high-RPE drift rate) had to be larger than β (the low-RPE drift rate) to create the aforementioned contextual “gap” after high-RPE items, which is how we explain impaired recognition priming in the high-RPE condition of Experiment 3, and impaired sequence memory in the high-RPE 1-trial-apart and 3-trial-apart conditions of Experiment 4.

The effects of γ^*FC*^ and γ^*CF*^ (episodic/semantic balance in the associative matrices) ended up being more complex. As discussed above, recognition priming depends on the semantic component of *M*^F C^ and the episodic component of *M*^*CF*^: The prime is loaded into context via *SM*^*F C*^, and then it cues the target via *EM*^*CF*^ (since the prime was part of the target’s episodic context during learning). Note that this is the same basic sequence of events that accounts for the forward bias in contiguity effects in free recall. The only difference is that, in free recall, the just-recalled item plays the role of the prime: the just-recalled item is loaded into context via *SM*^*F C*^ and cues recall of the following item via *EM*^*CF*^ (Howard & Kahana, 2002). Thus, to explain recognition priming effects (and forward contiguity effects in free recall), we need to ensure a substantial contribution of *SM*^*F C*^ and *EM*^*CF*^.

However, it would be unwise to fully “tilt” *M*^*F C*^ towards semantic memory and *M*^*CF*^ towards episodic memory. The episodic component of *M*^*F C*^ is also important: As noted earlier, this component is what gives rise to “mental time travel” effects in free recall -- in particular, backward transitions in free recall (i.e., recalling items that were studied *before* the item that was just recalled) are thought to result from a sequence where recalling an item reinstates that item’s context via *EM*^*F C*^, which then biases recall towards nearby items symmetrically in the backward and forward directions (Howard & Kahana, 2002). The semantic component of *M*^*CF*^ is also important: As described in *7.5*, our model uses this “direct readout” component to explain primacy effects in free recall -- the primacy item is (strongly) inserted into context via *SM*^*F C*^ and then is directly read out from context at test via *SM*^*CF*^. As an aside, this same mechanism that gives rise to primacy would also predict increased free recall of high RPE items (which, like primacy items, are assigned a higher-than-usual drift rate); we have not yet run an experiment to test this prediction in our paradigm.

To summarize the above: Both the episodic and semantic components of both *M*^*CF*^ and *M*^*F C*^ are important for explaining various effects (either effects in our data or classic regularities in free recall). As such, the greatest challenge in parameterizing the model was finding the right balance between the episodic and semantic components for each matrix. The fact that we found a set of parameters that works well for simulating our results (without impeding our ability to simulate primacy/recency/contiguity in free recall) serves as an existence proof that these factors *can* be suitably balanced.

## 8. General discussion

### 8.1. Summary of behavioral results

In a passive-viewing, Pavlovian reward learning task, we found that large reward prediction errors (RPEs) enhance memory and create event boundaries, thereby chunking rewarding experiences into discrete states in memory. Like other types of event boundaries, high RPEs enhance recognition for the event associated with the prediction error, while interrupting memory of the sequence of events around it. Specifically, we showed that high-RPE items demonstrate recognition priming, i.e, faster recognition of those items when primed by the previous item, indicating intact associative links with preceding events during encoding (Exp. 1-2). However, we found diminished recognition priming for events surrounding the high-RPE item (Exp. 3) providing evidence of an RPE-modulated event boundary. Moreover, we found that temporal memory, and in particular sequence memory, was worse for pairs that spanned a high RPE versus those that did not (Exp. 4). Interestingly, however, this worse overall sequence memory was only seen for pairs that did not include the high-RPE event itself, again showing that the high-RPE event is associated with its predecessor.

### 8.2. Summary of computational model

To illustrate and better understand the effects of event boundaries on memory in our experiments, we developed a computational model, a variant of the CMR model (Polyn et al., 2009), that qualitatively fits our results. To explain the effects of RPEs on memory, our model posits that large RPEs increase the drift rate of contextual information, effectively flushing out previous events and adding the current event into the drifting context.

We simulated recognition priming in the model and analyzed simulated recognition latencies; we also simulated the accuracy of sequential memory judgments. Using the mechanism described above (increased drift in response to large RPEs), we were able to explain our four most important experimental findings: 1) there was recognition priming for pairs of items that were presented sequentially at encoding, regardless of the size of the RPE associated with the target item; 2) when testing for priming of events that were separated by one event during encoding, recognition priming was disrupted if the intervening event triggered a high RPE; 3) when testing sequence memory for pairs of items that were presented adjacently at encoding, having one of those items be a high-RPE event did not impair sequence memory; 4) when testing sequence memory for pairs of items that were presented either one or three items apart at encoding, sequence memory was worse if that gap included a high-RPE event.

These simulation results illustrate the sufficiency of our drift-rate manipulation for explaining the effects of high (vs. low) RPE in the studies reported here. However, this demonstration of sufficiency does not rule out the possibility that RPEs can affect declarative memory in other ways. For example, in addition to (or instead of) increasing drift rate, RPEs might also increase the learning rate on item-context associations -- this would have the effect of stamping in the episodic memory of the high RPE event more strongly. More simulation work is needed to determine what combination of mechanisms does the best overall job of explaining the effects of RPEs on declarative memory.

### 8.3. High RPE events are better remembered

Consistent with previous results (Rouhani et al., 2018), we found that high RPEs led to better recognition memory for the event associated with the RPE. This finding is moreover consistent with work showing enhanced memory for other types of surprising events in the context of reward learning (Murty & Adcock, 2014; Murty, Labar, & Adcock, 2016), and outside of reward learning (Greve, Cooper, Kaula, Anderson, & Henson, 2017; Kalbe & Schwabe, 2019).

### 8.4. High RPEs form event boundaries in memory

We found that latent shifts in the reward value of a rewarding source induce event boundaries by interrupting the sequential integration of memories that occur before and after a high RPE, thus acting similarly to other event boundaries reported in the literature (DuBrow & Davachi, 2013, 2014; Ezzyat & Davachi, 2014; Horner et al., 2016). Heusser and colleagues (2018) recently demonstrated that enhanced associative memory for a perceptual boundary comes at the cost of integrating events across the boundary, reflecting a trade-off between the two processes. Here, we found concordant results in the domain of latent reward expectations: high-RPE events were not only better encoded but also demonstrated intact associative memory with their preceding items, through intact recognition priming and sequence memory. However, and in line with this trade-off, events surrounding the high RPE demonstrated diminished associative memory through impaired recognition priming and sequence memory.

We note that, in our task, we were interested in the effect of having detected a change in rewards (i.e., to induce sharp event boundaries), and so the jumps in the underlying reward distribution were quite obvious and sustained. In the real world, however, these changes may be more subtle and gradual, requiring multiple observations to infer an event boundary. Future work could introduce uncertainty around reward shifts and examine how this affects the temporal organization of events in memory (Dubrow et al., 2017).

Another key issue is whether the (apparent) contextual discontinuity evoked by high-RPE events in our study is attributable to the prediction error *per se*, or whether it is attributable to the fact that high RPEs indicated shifts in the underlying “latent cause” driving participants’ observations (see Zacks et al., 2007). In our paradigm, these two factors (RPE and shift-in-latent-cause) were confounded -- in future work, we can try to unconfound them (e.g, by having isolated high-reward or low-reward items that do not indicate a lasting change in the underlying mean reward value). Related to this point, Siefke et al. (2019) recently ran a study that attempted to unconfound context change and prediction error, using stimuli that varied in their background color; results from that study supported the hypothesis that context change, not prediction error *per se*, is the key determinant of discontinuities in mental context. More work is needed to see if this applies to our RPE paradigm.

### 8.5. Event boundary occurs across the high-RPE event

Although some theories (e.g., latent cause models, Gershman et al., 2014) predict that an event boundary occurs at the high prediction error event itself, separating that event from preceding items, we found intact associative links between the high-RPE event and its predecessor. At the same time, we found evidence for an event boundary across the high-RPE event. In our model, the high-RPE item and its predecessor are linked because the high-RPE −1 item is active in the context layer when the high-RPE item is presented at study. Additionally, the high-RPE item is strongly linked to its successor since the high-RPE item itself gets strongly inserted into the high-RPE +1 item’s context. For this reason, although we did not test for recognition priming between the high-RPE item and high-RPE +1 item, we predict, based on our model, that there will be strong recognition priming for the high-RPE +1 item when primed by the high-RPE item. Still, consistent with our behavioral results, the model predicts that recognition priming between the high-RPE −1 and the high-RPE +1 item will be disrupted because the increased drift associated with the high RPE leads to weak representation of the high-RPE −1 item in the high-RPE +1’s context. In sum, our model predicts that the high-RPE item is linked to both its predecessor and successor through context while disrupting the association of the events around it.

Previous work offers another potential mechanism for the preserved link between the high-RPE event and its preceding event, namely that at event boundaries, memory of the previous episode is reinstated (Sols et al., 2017), perhaps leading to binding between the high-RPE event and its predecessor. Other work has also shown that increased hippocampal activity at event offset (i.e., right after the boundary is inferred) predicts subsequent retrieval of the previous episode, in a sense “registering” the just-experienced episode (Baldassano et al., 2017; Ben-Yakov & Dudai, 2011; Ben-Yakov et al., 2013). In our task, the boundary itself is calculated by the difference between the expected value and the current reward, which, along with the “replay” mechanism described above, could additionally bind the high-RPE event with its predecessor.

### 8.6. Recognition priming for high-versus low-RPE events

Our results in Experiments 1 and 2 were suggestive of stronger recognition priming for high-RPE items than for low-RPE items (i.e., numerically, high-RPE items were retrieved faster than the low-RPE targets, although not significantly). We note that in previous studies, recognition priming was evident only for high-confidence recognition (i.e., for recollection instead of familiarity; DuBrow & Davachi, 2013; Schwartz et al., 2005), and we did not collect confidence judgments in our task, perhaps occluding more stable recognition priming effects in the low-RPE pairs. If anything, however, this emphasizes the intact association of the high-RPE event with its predecessor, as we saw recognition priming for the high-RPE item across all confidence levels.

### 8.7. Distance memory

Event boundaries increase the subjective temporal distance between events (Ezzyat & Davachi, 2014). We saw this effect only in the replication of Experiment 4, which points to more variable results with this measure. Across both datasets in Experiment 4, participant response was quite inaccurate and was in fact highest for the average value of the response scale (“3 trials apart”), even though two-thirds of the actual distances smaller than 3. Nevertheless, the greater subjective distance for high-RPE pairs was largely driven by the pair that included the high-RPE event itself (“0 trials apart”). However, this pair did not demonstrate impaired sequence memory, which points to perhaps a dissociation between mechanisms supporting sequence and distance memory.

### 8.8. Neural mechanisms

RPEs modulate dopamine release in the ventral tegmental area (VTA) by increasing firing when rewards are better than expected, and decreasing firing when rewards are worse than expected (Barto, 1995; Montague, Dayan, & Sejnowski, 1996). Given dopamine-dependent plasticity in the hippocampus, associated with memory formation, putative links have been made between RPE signals in the VTA and modulation of hippocampal plasticity (Lisman & Grace, 2005), giving rise to enhanced memory for events that are better than expected (Jang, Nassar, Dillon, & Frank, 2019).

Recent work also offers a mechanism by which unsigned (absolute value) RPEs interact with memory. The locus coeruleus (LC), a previously unknown source of dopamine, co-releases dopamine along with its known release of norepinephrine, to generate hippocampal memories (Kempadoo, Mosharov, Choi, Sulzer, & Kandel, 2016; Takeuchi et al., 2016). Large RPEs, whether positive or negative, have been shown to increase learning rate during reward learning, and are thought to modulate the noradrenergic LC system and its connections to the anterior cingulate cortex (Behrens, Woolrich, Walton, & Rushworth, 2007; Courville, Daw, & Touretzky, 2006; Nassar et al., 2012; Roesch, Esber, Li, Daw, & Schoenbaum, 2012; Sara, 2009) -- a system linked to memory for surprising or arousing events (Clewett, Huang, Velasco, Lee, & Mather, 2018; Clewett, Schoeke, & Mather, 2014). Moreover, prediction errors are thought to enact a “network reset” (Zacks et al., 2007) that has been recently linked to a shifting latent-state representation in the orbitofrontal cortex (Nassar, McGuire, Ritz, & Kable, 2018). The orbitofrontal cortex is a strong candidate region for representing these latent states (Schuck, Cai, Wilson, & Niv, 2016), which are thought to encode a cognitive map of task space (Wilson, Takahashi, Schoenbaum, & Niv, 2014). Seeing that event boundaries modulate representations in the hippocampus (DuBrow & Davachi, 2014; Ezzyat & Davachi, 2014), it has been suggested that at these boundaries, enhanced hippocampal activity and a shift in cortical representations (such as in the orbitofrontal cortex) increases the drift in temporal context (Brunec, Moscovitch, & Barense, 2018). Future work should characterize the interactions between the orbitofrontal cortex and the hippocampus in segmenting our experiences and organizing those memories.

## 9. Conclusion

Across four experiments, we found that latent shifts in the mean value of a reward distribution, generating the experience of high reward prediction errors, led to stronger recognition for the event associated with the high prediction error, but interrupted the sequential integration of events around the prediction error, thereby acting like an event boundary in memory. We developed a computational model that treats a high prediction error event as an increase in the updating of that event to an internal, temporal context during encoding (thus creating a representational break between the events that occurred before and after the high prediction error event), and were able to capture our recognition priming and sequence memory results. These results suggest that large changes in the value of a rewarding experience split our memories of those experiences, separating them into separate clusters in memory, each including similarly rewarding events. This mechanism can help create low-dimensional representations of task states that are useful for both learning and decision making.

## Acknowledgements

We thank Sarah Dubrow for her instrumental consulting on this project, and Lynn Lohnas, Per Sederberg, and Rivka Cohen for fruitful conversation. This work was supported by grant W911NF-14-1-0101 from the Army Research Office (Y.N.), grant R01MH098861 from the National Institute for Mental Health (Y.N.), and the National Science Foundation’s Graduate Research Fellowship Program (N.R.).

## Supplemental Material

**Table 1.**
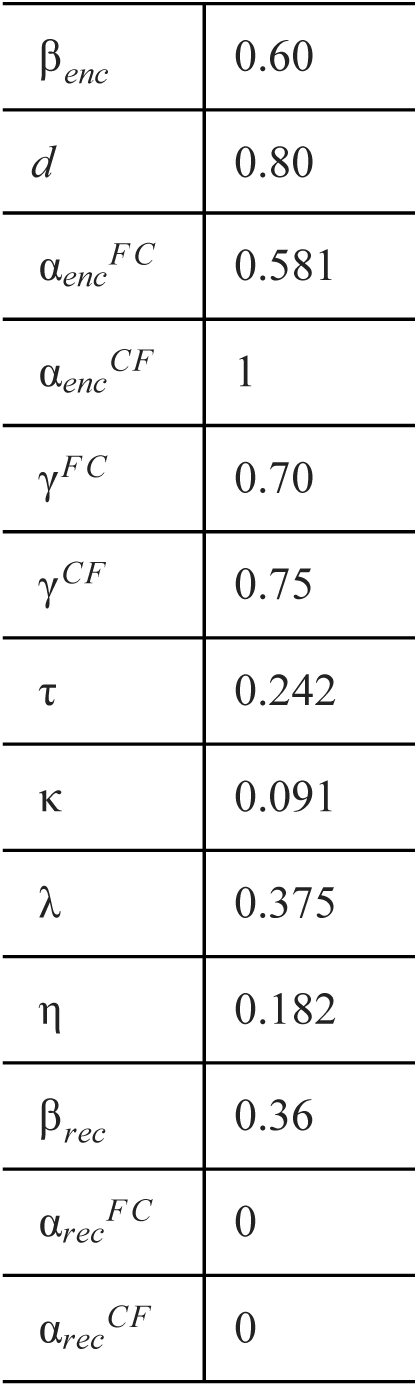
Parameters used in model. β_*enc*_, *d*, γ^*FC*^, and γ^*CF*^ were determined based on a parameter search for values that best captured primacy and recency effects in serial position curves, as well as the signature characteristics of conditional response probability curves in free recall. All other parameters were taken from Polyn et al. (2009); “*enc*” refers to the value used during encoding and “*rec*” during recall.

|  |  |
| --- | --- |
| $\beta_{enc}$ | 0.60 |
| $d$ | 0.80 |
| $\alpha_{enc}^{FC}$ | 0.581 |
| $\alpha_{enc}^{CF}$ | 1 |
| $\gamma^{FC}$ | 0.70 |
| $\gamma^{CF}$ | 0.75 |
| $\tau$ | 0.242 |
| $\kappa$ | 0.091 |
| $\lambda$ | 0.375 |
| $\eta$ | 0.182 |
| $\beta_{rec}$ | 0.36 |
| $\alpha_{rec}^{FC}$ | 0 |
| $\alpha_{rec}^{CF}$ | 0 |

